## Supplemental figures and tables for "Genome-centric analysis of short and long read metagenomes reveals uncharacterized microbiome diversity in Southeast Asians"

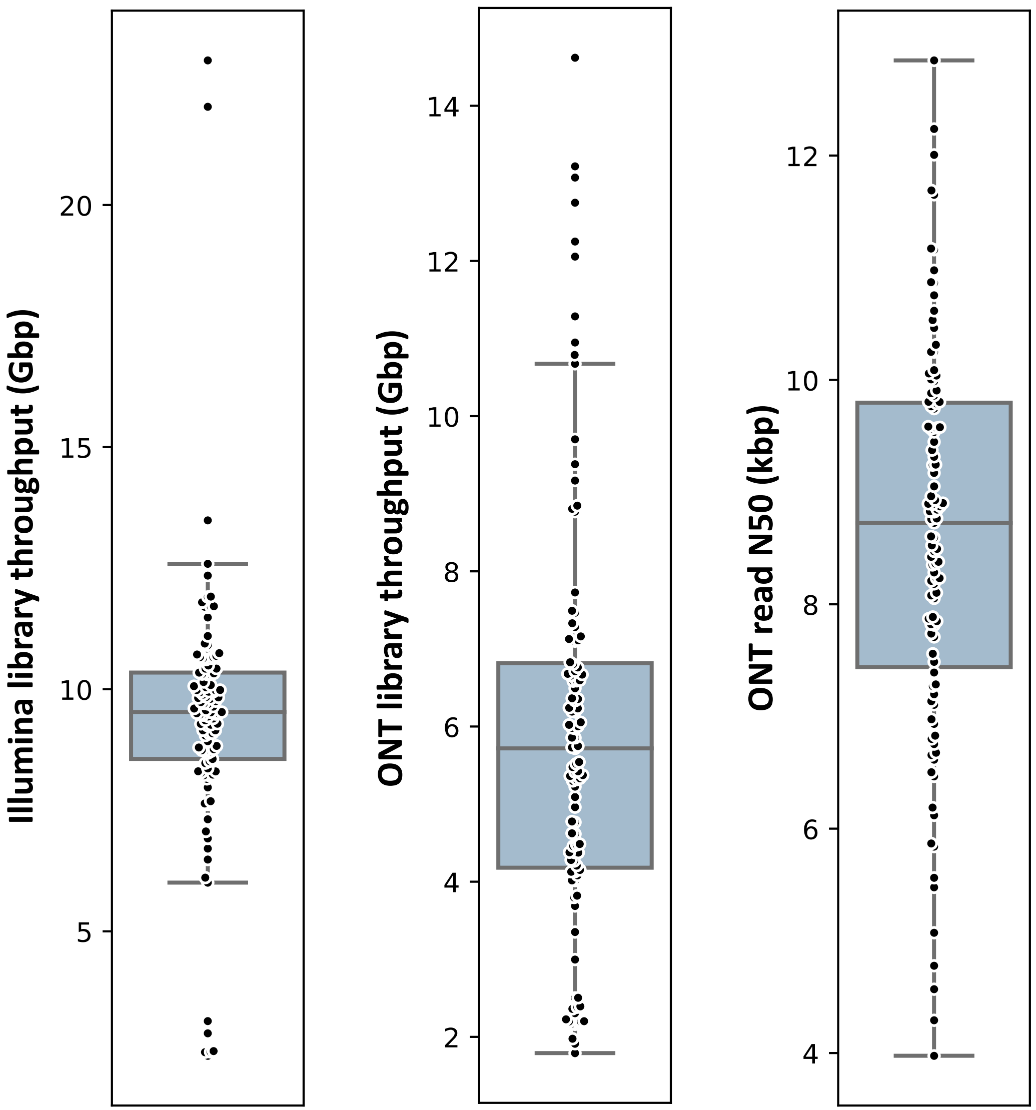


**Supplementary Figure 1**: **Long and short read sequencing library statistics.** Boxplots depicting the distribution of sequencing throughput for Illumina and ONT libraries, as well as ONT read N50 (read length such that >50% of the sequence data is in longer reads), across samples in SPMP (n=109). Center lines in the boxplots represent median values, box limits represent upper and lower quartile values, whiskers represent 1.5 times the interquartile range above the upper quartile and below the lower quartile, and all data points are represented as dots in the figure.

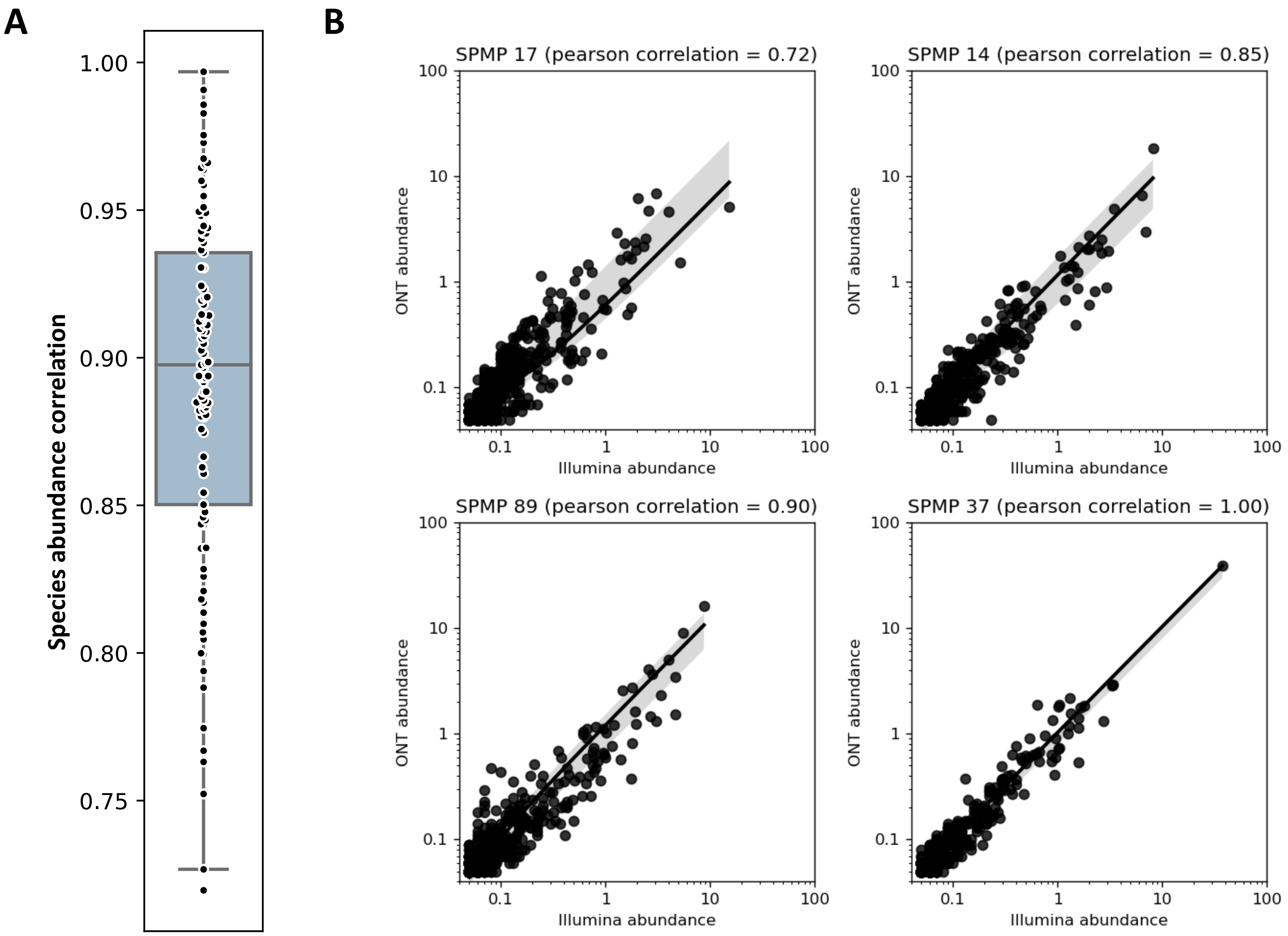


**Supplementary Figure 2**: **Concordance in species composition across sequencing technologies.** (A) Boxplot showing the distribution of Pearson correlation values obtained when comparing species-level relative abundance profiles with Illumina and ONT reads for SPMP samples (n=109). Center line represents median value, box limits represent upper and lower quartile values, whiskers represent 1.5 times the interquartile range above the upper quartile and below the lower quartile, and all data points are represented as dots in the figure. Relative abundances were obtained from Bracken and Kraken2 analysis using the UHGG database. (B) Scatter-plots showing the relative abundances (log scale) for different species obtained using Illumina reads (x-axis) and ONT reads (y-axis) for a selected set of 4 samples that span the range of correlation values.

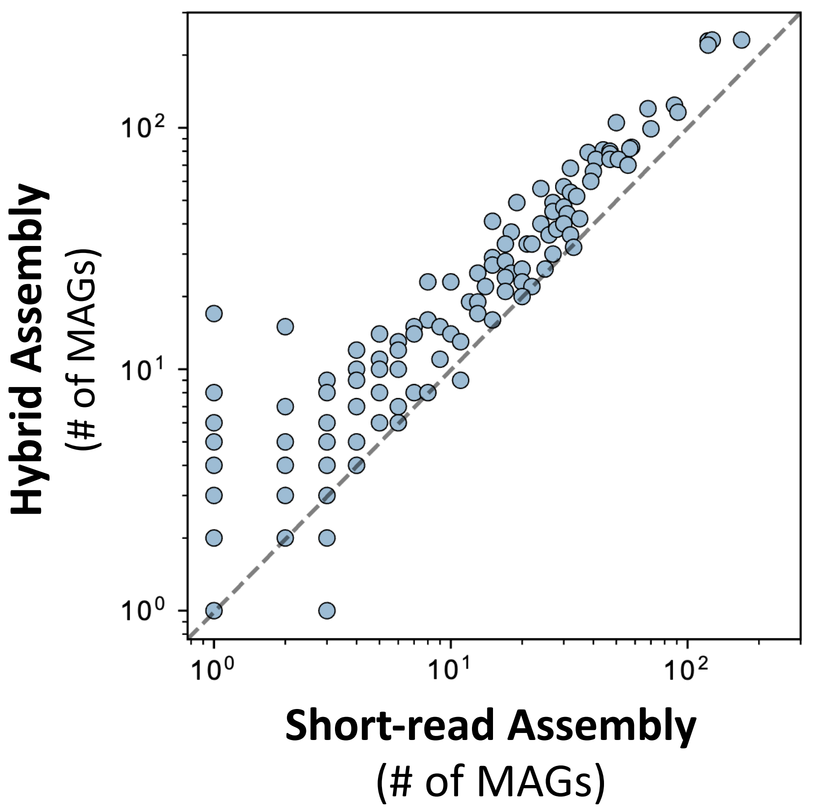


**Supplementary figure 3**: **Genus-wide comparison of MAGs obtained with short-read versus hybrid assembly.** Scatterplot showing the number of MAGs obtained for each genus from short-read versus hybrid assembly. The diagonal line represents a ratio of 1 between the two datasets. No bias was detected towards any genera (Fisher's exact p-value>0.05, Bonferroni corrected).

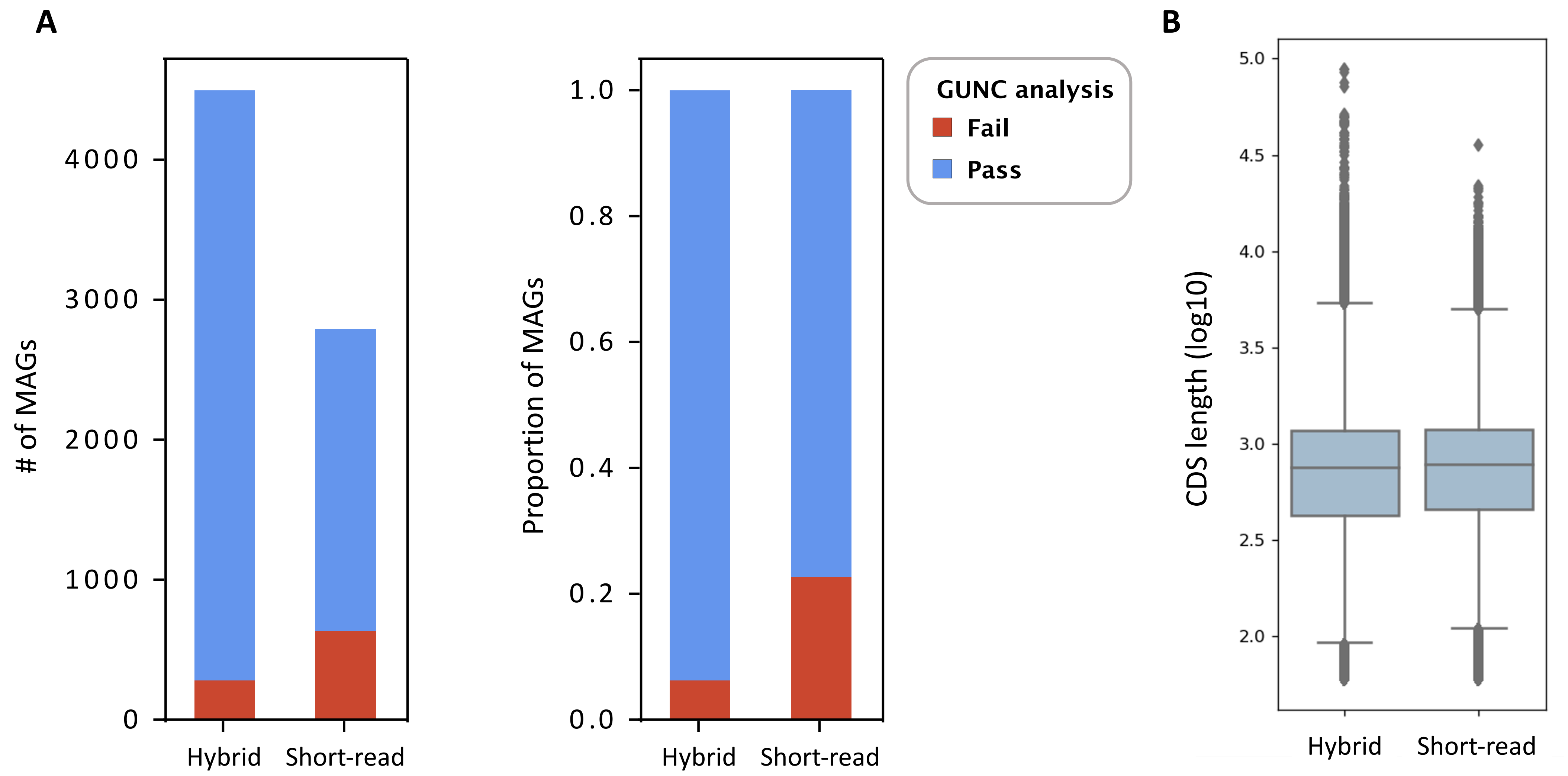


**Supplementary figure 4**: **Assessment of MAG quality in terms of chimerism and gene lengths.** (A) Barplots showing the number and the proportion of MAGs passing or failing chimerism detection analysis with GUNC. (B) Boxplots showing coding sequence (CDS) length distributions for short-read and hybrid assemblies based on Prodigal gene predictions. The distributions are similar in both cases, with median lengths of 750bp and 783bp for hybrid and short-read assemblies, respectively, suggesting that indel errors may not be significantly impacting the ability to call genes with hybrid assemblies.

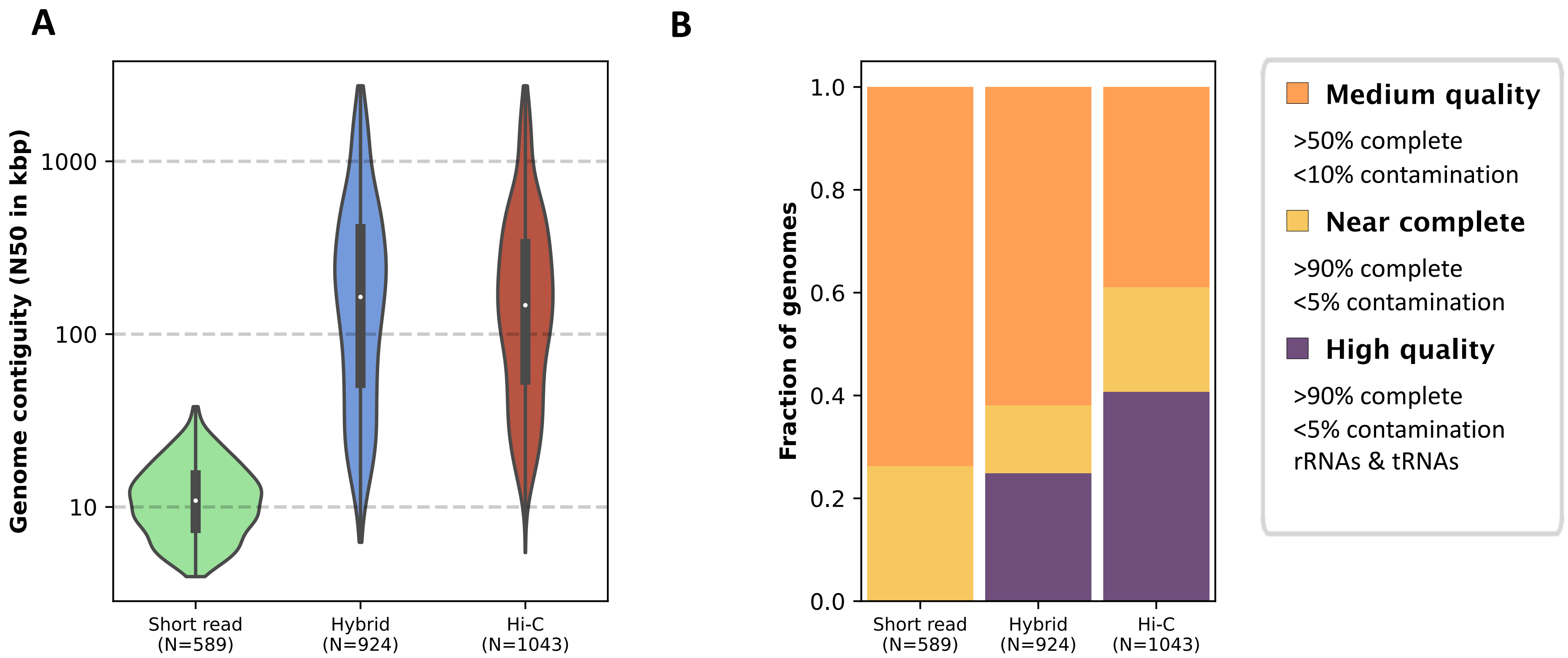


**Supplementary figure 5: Comparison of assembly contiguity and quality using Hi-C data.** Results reported here are an extension of those in **Figure 1D, 1E** where the hybrid assemblies were further scaffolded with Hi-C data (n=24). (A) Violin plots showing the distribution of N50 values obtained for MAGs generated with *short read* data, using a *hybrid* assembly, and after augmenting with *Hi-C* data. (B) Quality of corresponding MAGs based on MIMAG criteria.

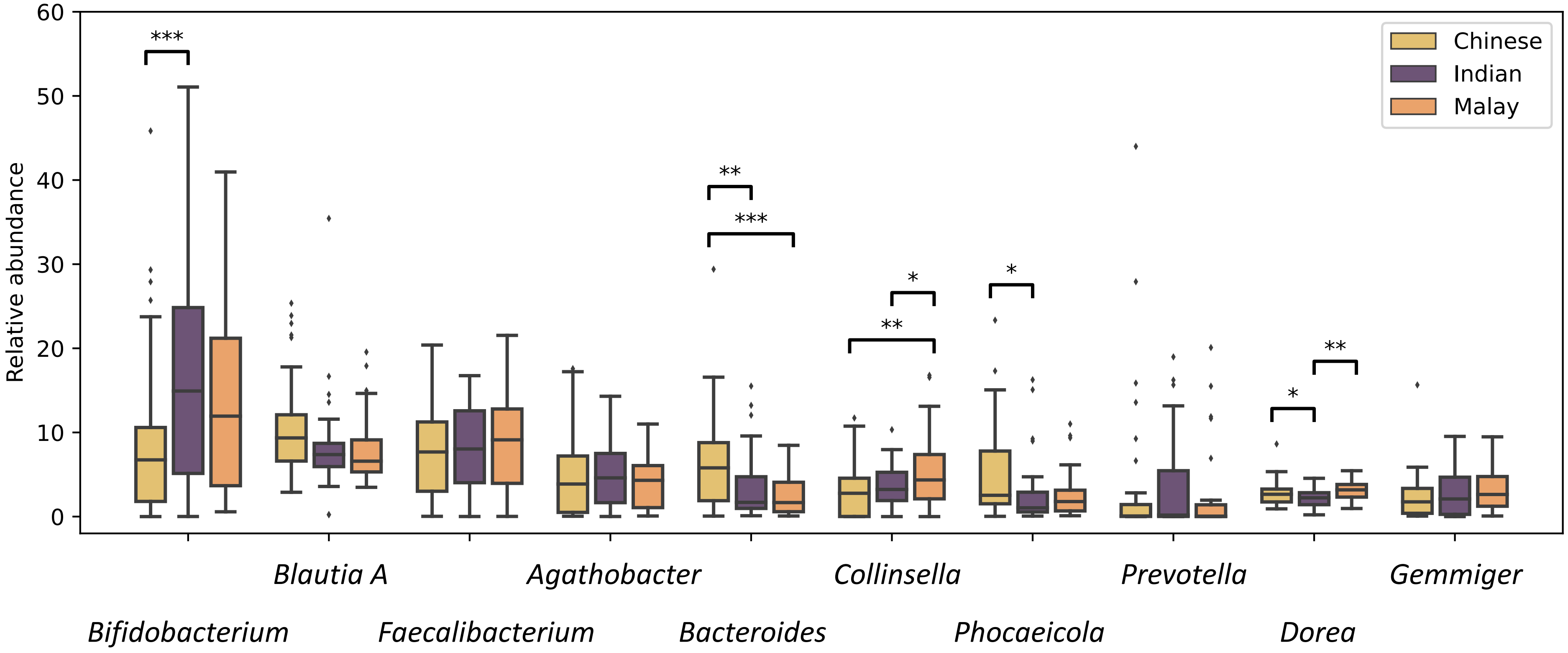


**Supplementary figure 6: Variability in relative abundance of common gut bacterial genera across ethnic populations in Singapore.** Boxplots depicting distribution of relative abundances for various gut bacterial genera across ethnicities (Chinese: n=53, Indian: n=30, Malay: n=26). The top 10 most abundant genera were selected for visualization here (median abundance across all subjects). Center line represents median value, box limits represent upper and lower quartile values, whiskers represent 1.5 times the interquartile range above the upper quartile and below the lower quartile, and all data points are represented as dots in the figure. Significant differences are indicated with stars (MaAsLin2 p-value<0.05: “*”, p-value<0.01: “**”, p-value<0.001: “***”).

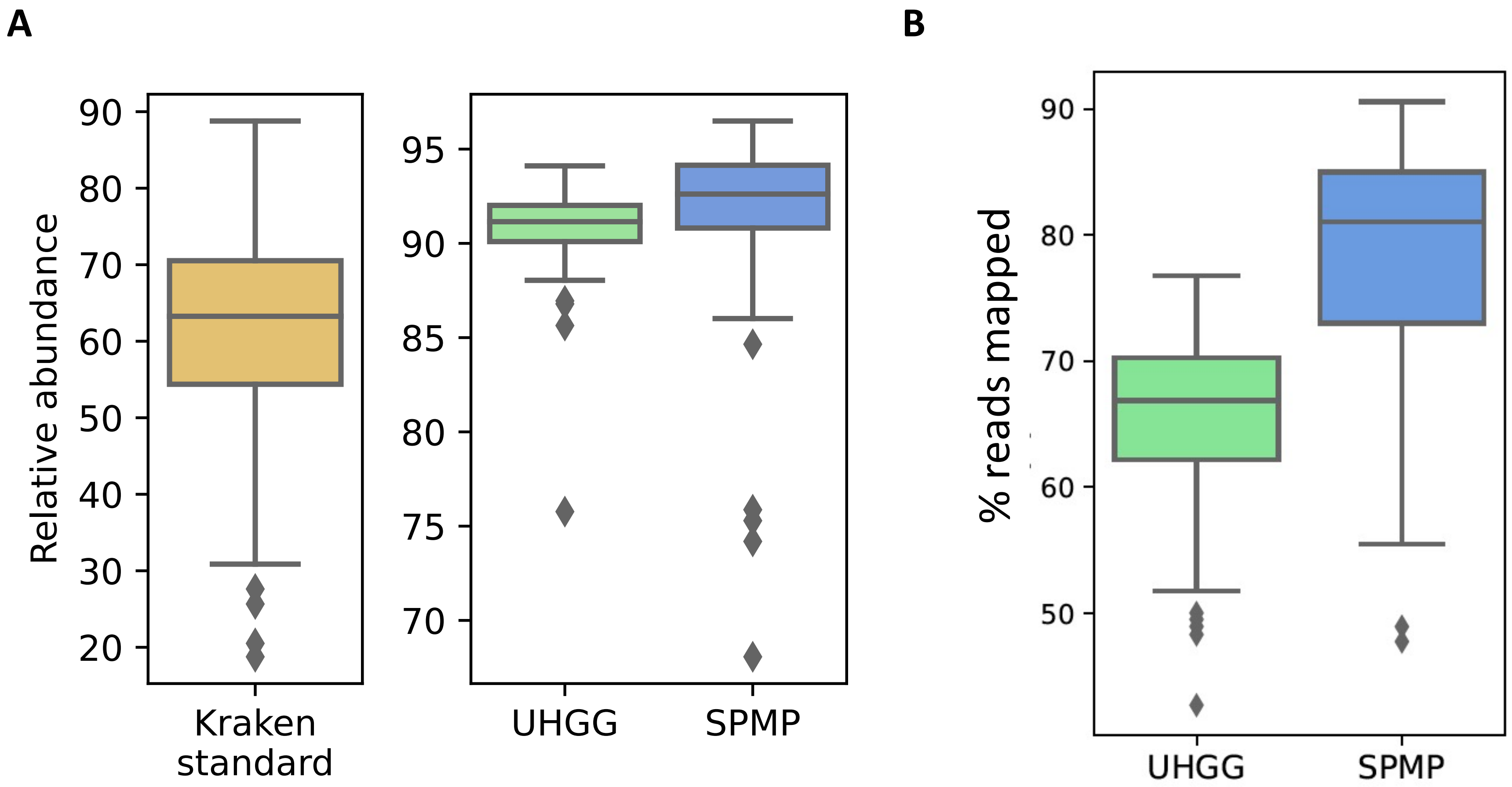


**Supplementary figure 7**: **Utility of SPMP database for gut bacterial read classification and mapping.** (A) Boxplots showing the percentage of gut bacterial reads identified using the standard database for Kraken, the UHGG database and SPMP references. Reads were obtained for an independent collection of 144 gut microbiomes from a clinical study in Singapore (manuscript under review). (B) Boxplots showing the percentage of reads mapped to UHGG (n=4,644, dereplicated database) and SPMP genomes (n=4,497). Note that both databases have similar sizes, while the full UHGG database is much larger, with a memory footprint that makes such analysis infeasible (>50×). Reads were mapped with minimap2, allowing for 2 or fewer mismatches and minimum alignment length of 100bp. Center line represents median value, box limits represent upper and lower quartile values, whiskers represent 1.5 times the interquartile range above the upper quartile and below the lower quartile, and all data points are represented as dots in the figure.

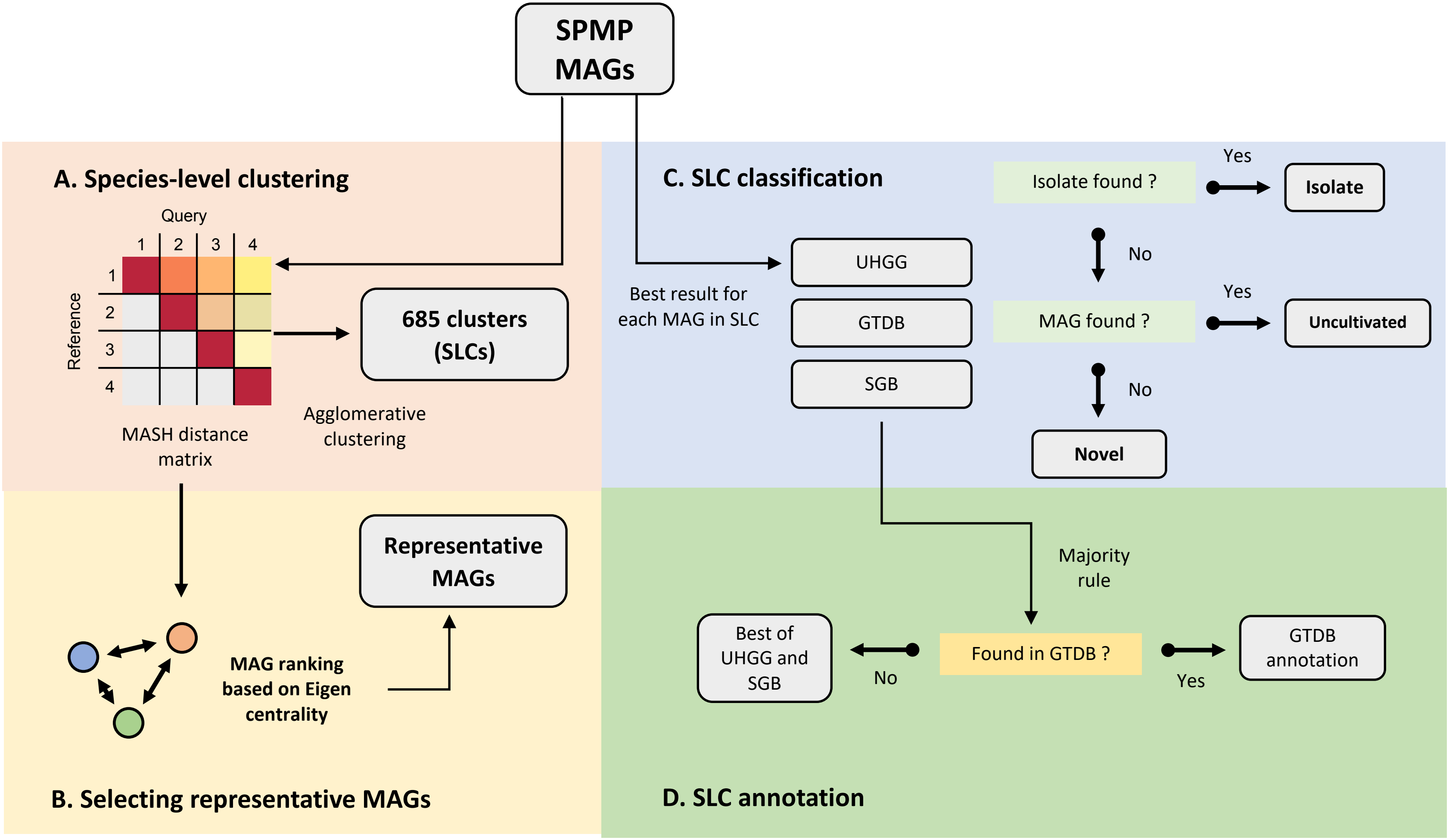


**Supplementary figure 8**: **Schematic representation of workflow for obtaining species-level clusters and annotating them with genomic databases.** Species-level clusters (SLCs) were obtained from MAGs based on pairwise MASH distances and an Agglomerative clustering approach (subfigure A). For each species-level cluster, a weighted graph was produced using mash distances to define representative MAGs based on Eigen centrality (NetworkX; subfigure B). SLCs were then annotated based on the best MASH result of each SLC's MAGs against 3 major databases, into those that have *isolate* strains, are composed solely of known MAGs (*uncultivated*) or are *novel* in SPMP (subfigure C). Finally, using all mash results, SLCs were annotated with taxonomic IDs based on existing databases (subfigure D).

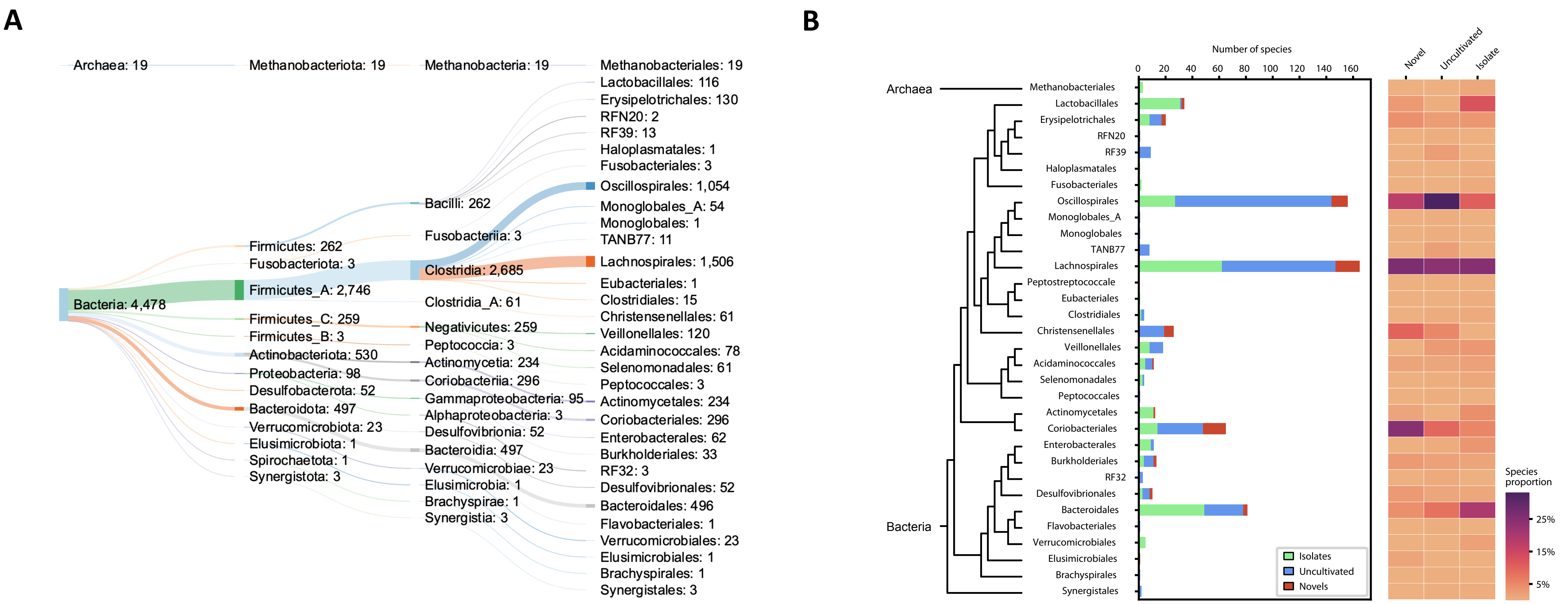


**Supplementary figure 9**: **Phylogenetic distribution of MAGs and species-level clusters in SPMP.** (A) Sankey plot showing the distribution of MAGs in different phylogenetic groups (GTDBtk assignment to Division, Phylum, Class and Order, from left to right; [http://sankeymatic.com](http://sankeymatic.com)) and (B) Distribution of species-level clusters shown with the order level phylogenetic tree (left), stacked barchart showing breakdown in terms of isolate, uncultivated and novel SLCs (middle), and heatmap showing corresponding relative proportions (right).

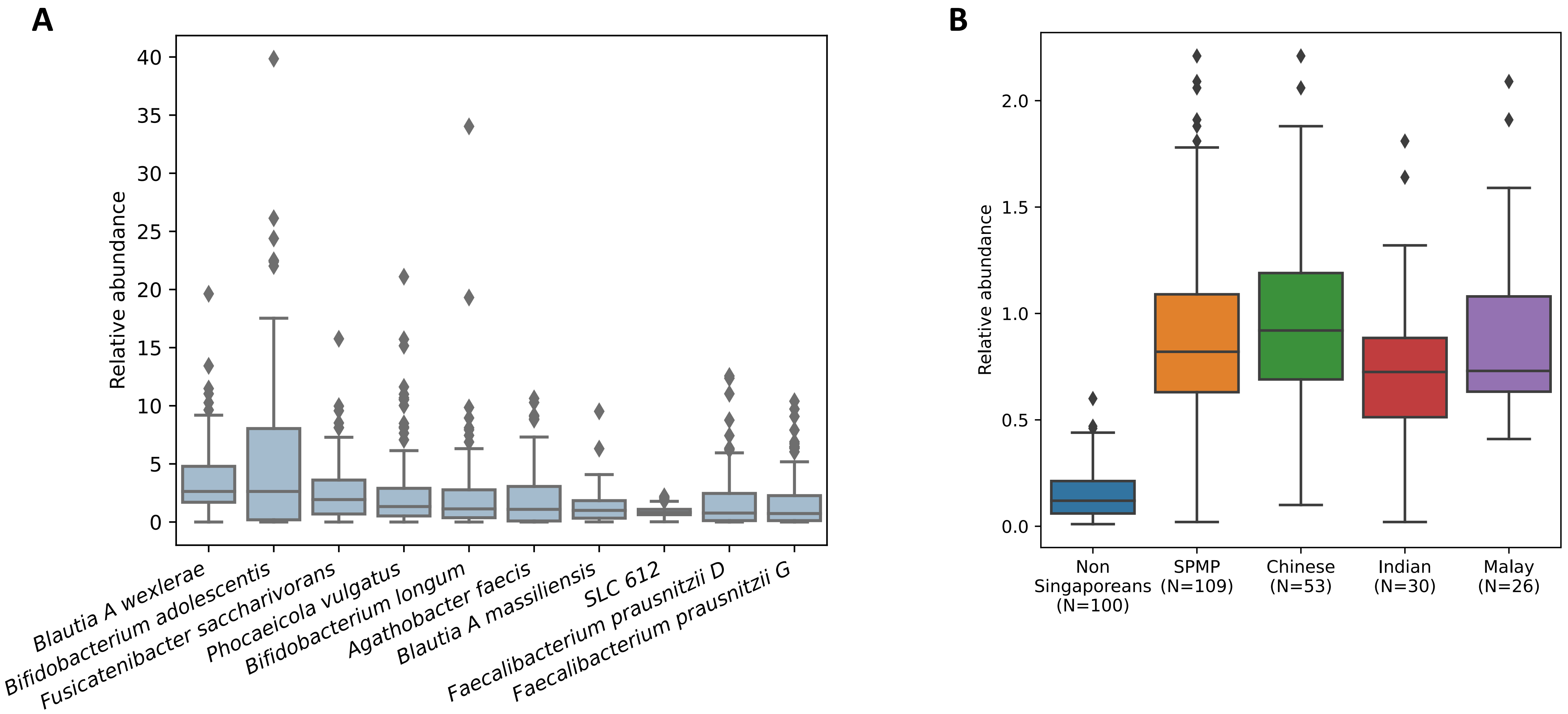


**Supplementary figure 10**: (A) Top 10 most abundant species in the gut microbiomes of SPMP subjects. (B) Relative abundance of SLC612 in the gut microbiomes of SPMP subjects versus non-Singaporean subjects (HMP). Center lines represents median value, box limits represent upper and lower quartile values, whiskers represent 1.5 times the interquartile range above the upper quartile and below the lower quartile, and all data points are represented as dots in the figures.

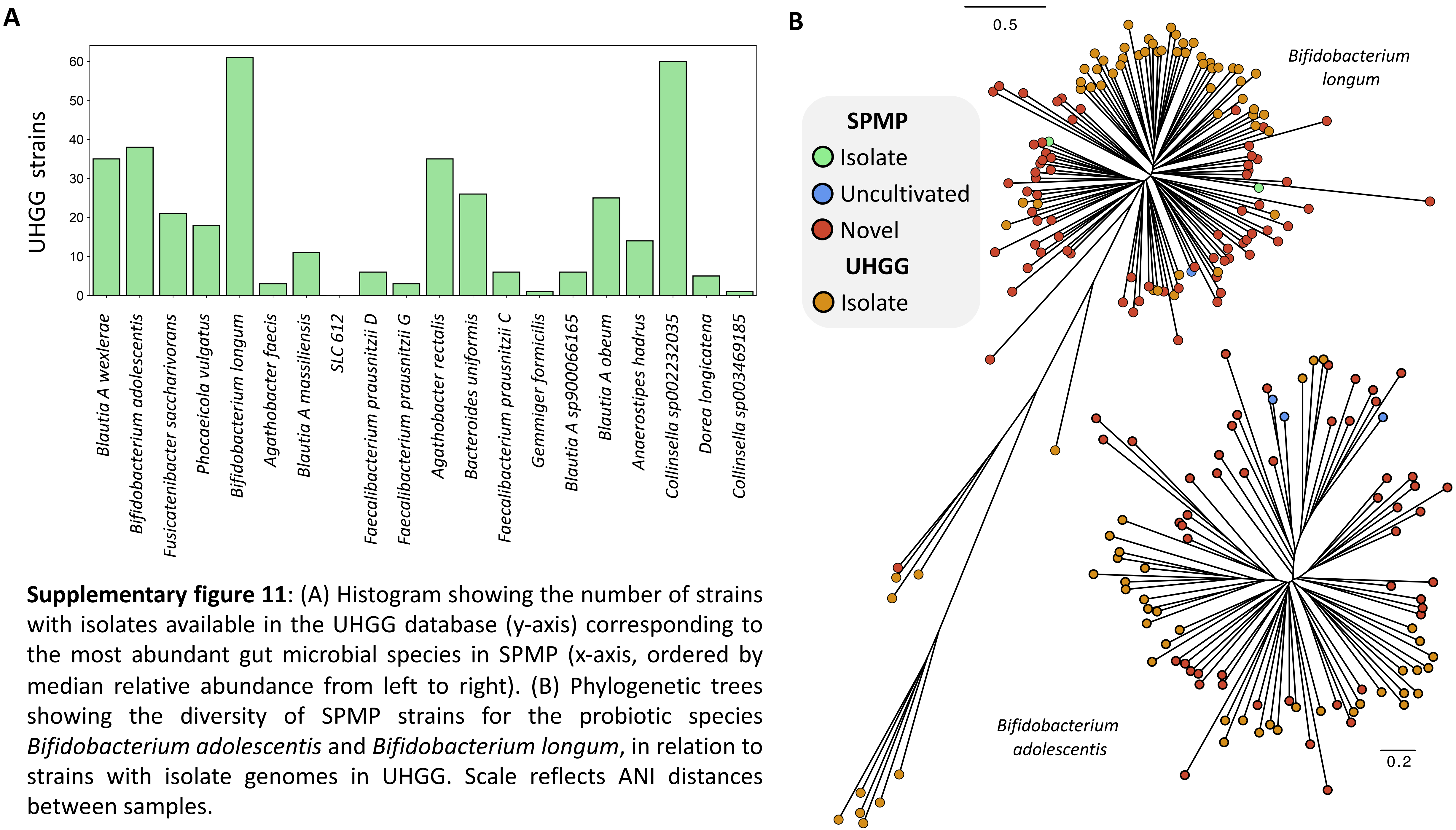


**Supplementary figure 11**: (A) Histogram showing the number of strains with isolates available in the UHGG database (y-axis) corresponding to the most abundant gut microbial species in SPMP (x-axis, ordered by median relative abundance from left to right). (B) Phylogenetic trees showing the diversity of SPMP strains for the probiotic species *Bifidobacterium adolescentis* and *Bifidobacterium longum*, in relation to strains with isolate genomes in UHGG. Scale reflects ANI distances between samples.

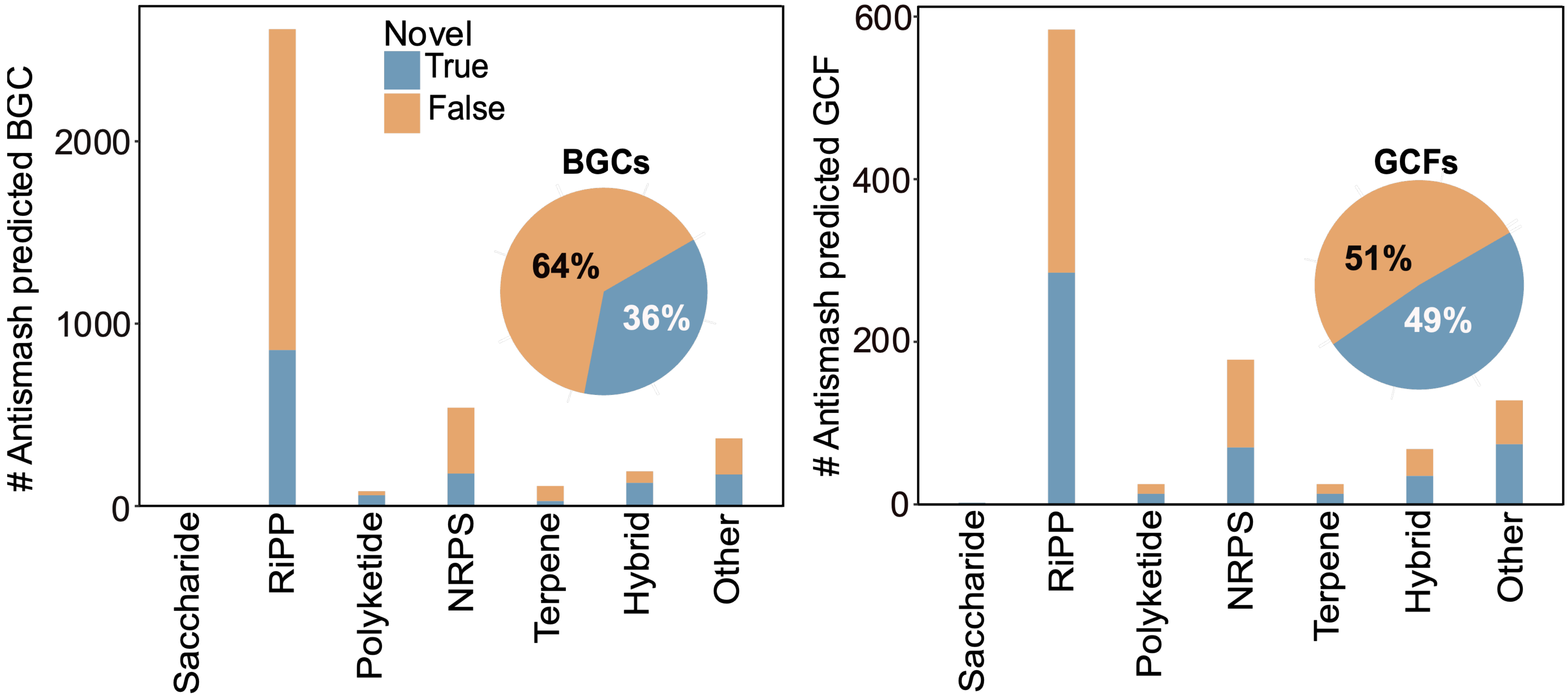


**Supplementary figure 12: Many SPMP BGCs predicted by antiSMASH are novel.** Stacked barcharts showing the number of SPMP antiSMASH predicted BGCs (left) and GCFs (right) in different product classes that are either present in annotated BGCs from HRGM and the antiSMASH/MIBiG databases, or absent. The latter group of SPMP BGCs are considered novel. Inset piecharts show the overall breakdown.

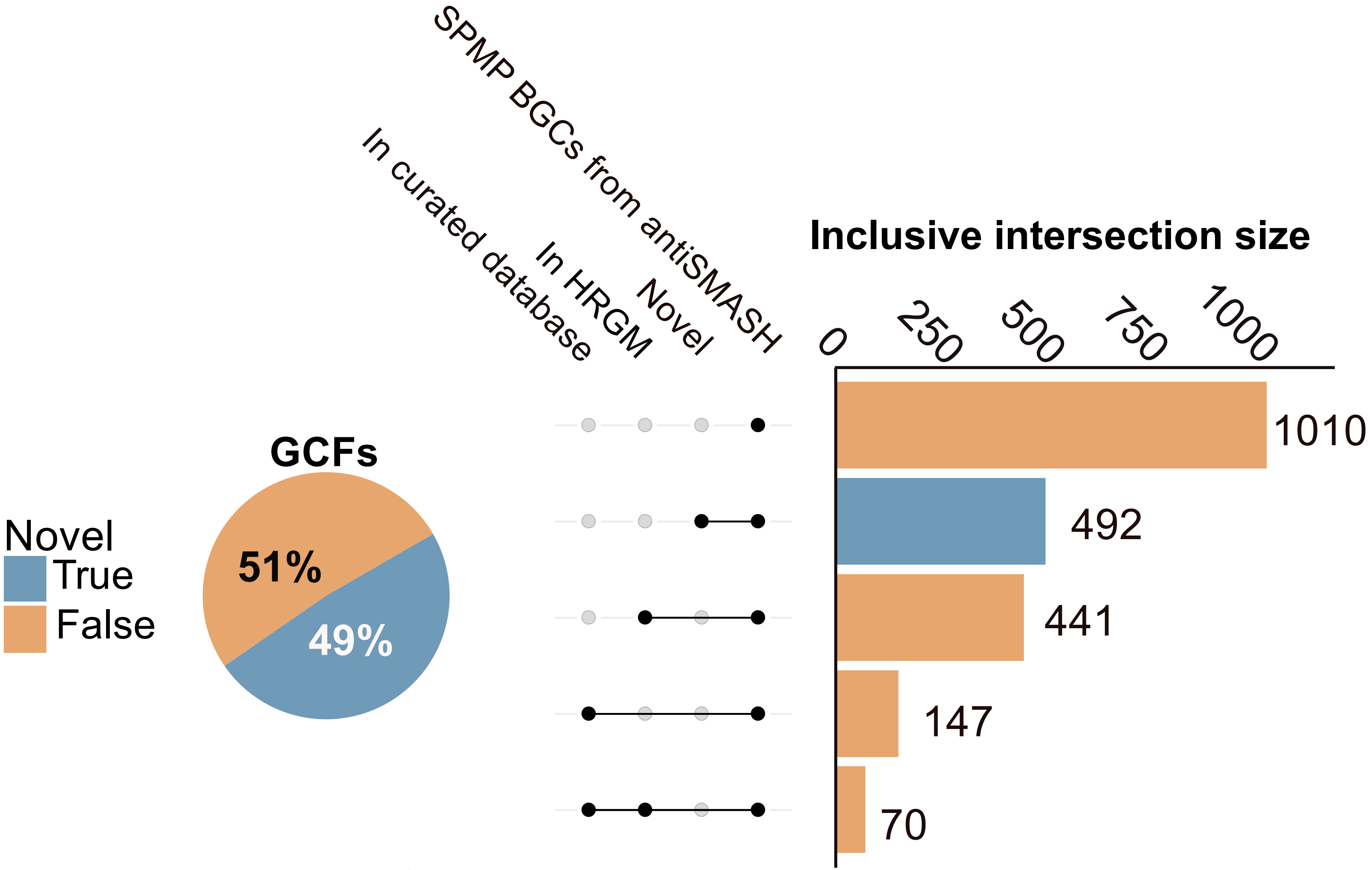


**Supplementary figure 13: Distribution of SPMP BGCs predicted by antiSMASH in different categories based on similarities to existing annotations or novelty.** Upset plot showing overlaps in SPMP GCFs predicted by antiSMASH with sequences in curated BGC databases or in HRGM antiSMASH annotations. Numbers represent the size of the intersection between different combinations of sets. The piechart shows the overall breakdown.

| Unified Label | DeepBGC Labels | antiSMASH Labels |
|---|---|---|
| RiPP | RiPP | lanthipeptide; bacteriocin; thiopeptide; lassopeptide; cyanobactin; sactipeptide; linaridin; bottromycin; microviridin; head_to_tail; glycocin; LAP; lipolanthine; proteusin; microcin; RaS-RiPP |
| NRPS | NRP | NRPS; NRPS-like |
| Polyketide | Polyketide | T1PKS; T3PKS; transAT-PKS; hglE-KS |
| Saccharide | Saccharide | oligosaccharide; amglyccycl |
| Others | Terpene; Others | terpene; arylpolyene; betalactone; TfuA-related; butyrolactone; ladderane; CDPS; siderophore; phenazine; resorcinol; phosphonate |

**Supplementary table 1**: Mapping of biosynthetic gene cluster class labels from DeepBGC and Antismash into a unified set of labels in this study.
